# Gastrodin ameliorates synaptic impairment, reestablishes mitochondrial membrane potential and reduces oxidative stress in N2a/APP cells through ERK1/2 and GSK-3β pathways

**DOI:** 10.1101/2023.01.15.524095

**Authors:** Zhi Tang, Yaqian Peng, Li Wang, Min Guo, Zhuyi Chen, Ting Zhang, Yan Xiao, Ruiqing Ni, Xiaolan Qi

## Abstract

Alzheimer’s disease (AD) is featured by abnormal β-amyloid (Aβ) deposition, neurofibrillary tangle formation, downstream mitochondrial dysfunction, oxidative stress, and synaptic loss. Gastrodin, a phenolic glycoside, has shown neuroprotective effect and used in the treatment of a range of brain diseases. Here we aim to assess the mechanisms and signaling pathways involved in the neuroprotective effect of gastrodin in murine neuroblastoma N2a cells expressing human Swedish mutant amyloid precursor protein (N2a/APP). The levels of pre- and postsynaptic proteins, amyloid precursor protein C-terminal fragments (APP-CTFs), levels of tau, glycogen synthase kinase-3 β (GSK-3β), extracellular regulated kinase (ERK), and c-Jun N-terminal Kinase (JNK) were assessed by Western blotting. Flow cytometry assays for mitochondrial membrane potential (JC1) and reactive oxidative stress, as well as immunofluorescence staining for lipid peroxidation (4◻hydroxynonenal) and DNA oxidation (8◻hydroxy◻2’◻deoxyguanosine), were performed. We found that gastrodin treatment increased the levels of presynaptic SNAP25, synaptophysin, and postsynaptic PSD95, reduced phosphorylated tau protein Ser396, and APP-CTFs in N2a/APP cells. In addition, gastrodin reduced the levels of reactive oxygen species generation, lipid peroxidation, and DNA oxidation, reestablished mitochondrial membrane potential. Upregulated levels of phosphorylated-GSK-3β, reduced levels of phosphorylated-ERK, and phosphorylated-JNK were involved the protective effect of gastrodin. In conclusion, we demonstrated a neuroprotective effect of gastrodin in N2a/APP cell line.

## 1. Introduction

Alzheimer’s disease (AD) is pathologically characterized by extracellular deposition of amyloid-beta (Aβ) and intracellular formation of neurofibrillary tangles associated with loss of synapses and neurons [1]. Amyloid precursor protein (APP) is cleaved by β-site APP-cleaving enzyme 1 (BACE1) in the amyloidogenic pathway producing Aβ [2, 3]. Aβ plays a crucial role in the pathogenesis of AD [4–7]. Synergistic interaction between Aβ and tau leads to downstream oxidative stress, gliosis, synaptic dysfunction, and neurodegeneration [8]. In addition, the accumulation of Aβ and p-tau has been shown to contribute to mitochondrial structural and functional changes [9–12] The exact molecular mechanisms underlying AD are not fully clear. Oxidative stress plays an essential role in many neurodegenerative diseases, including diabetes, and is upregulated in AD [13–17] Earlier studies have shown that Aβ accumulation [5–7, 18] and abnormal tau hyperphosphorylation [19–21] induce increased oxidative stress and reactive oxygen species (ROS) production, leading to synaptic impairment and neurodegeneration [13, 22, 23].

Gastrodin, 4-hydroxybenzyl alcohol-4-O-β-D-glucopyranoside, is a bioactive phenolic glycoside extracted from the orchid Gastrodia elata and Rhizoma Gastrodiae [24]. Gastrodin has been used in traditional medicine and exerts many pharmacological activities, including antidepressant, anti-inflammatory, and neuroprotective effects [24–31] Emerging evidence has shown that gastrodin improves cognitive decline in mouse models of AD, Parkinson’s disease, and vascular dementia [32–36] Multiple modes of action of gastrodin in AD cellular and animal models have been demonstrated [24], such as suppressing BACE1 expression [35, 37], reducing tau hyperphosphorylation and Aβ aggregation, and blocking Aβ-induced cytotoxicity in a cell model and cognitive impairment in animal models [35, 38, 39]. Gastrodin treatment has been shown to decrease Aβ-induced cell death and decrease lactic dehydrogenase release in primary cultured neuronal cells [40]. In addition, gastrodin exerts antioxidant, mitochondrial protection [41, 42] and anti-inflammatory activities [32, 34, 41–45]. Several signaling pathways have been found mediating the effect of gastrodin in these studies, including nuclear factor-E2-related factor 2 (Nrf2), extracellular regulated kinase (ERK) 1/2, mitogen activated protein kinase (MAPK) and glycogen synthase kinase-3β (GSK-3β) [26, 28, 34, 37, 43, 46–50].

Here, we examined the effect of gastrodin on ameliorating synaptic impairment, APP processing, tau phosphorylation, mitochondrial dysfunction, and oxidative stress damage in a cell model of AD (N2a/APP). The GSK-3β or ERK1/2 pathway is one of the most important signaling pathways in the central nervous system [51–54] We hypothesized that the ERK1/2, GSK-3β and c-Jun N-terminal kinase (JNK) signaling pathways may be involved in the effect of gastrodin.

## 2. Methods and Materials

### 2.1 Reagents, antibodies, cell culture and treatment

The list and sources of materials, including chemicals, antibodies, and kits, are described in detail in **Supplemental Tables 1** and **2**. Gastrodin (purity ≥98.0%) was dissolved first in dimethyl sulfoxide (DMSO, 10% v/v) and then further diluted with polyethylene glycol 300 (PEG300, 40%), Tween-80 (5%) and saline (45%). N2a/APP cells stably transfected with the human APP Swedish mutant (N2a/APP) were gifts from Prof. Rong Liu (Tongji Medical School, Wuhan, China). Cells were cultured to 90% confluence in a T25 culture flask in Dulbecco’s modified Eagle’s medium (DMEM) with 10% (v/v) fetal bovine serum (FBS) and 1% (v/v) penicillin/streptomycin solution in an incubator under 5% CO2 at 37 °C. At 80-90% confluence, cells were seeded into 6-well culture plates. N2a/APP cells were treated with different concentrations of gastrodin (1 μM, 5 μM, and 10 μM) for 24 h in an incubator.

### 2.3 Cell viability experiments

Cell viability was examined using Cell Counting Kit-8 (CCK-8) as previously described [55]. Briefly, N2a/APP cells were seeded in 96-well plates at nearly 5000 cells per well. After gastrodin treatment, 10 μL of CCK-8 reagent and 100 μL of medium were added to each well and incubated for 1 h at 37°C. After incubation, the absorbance of the plate was measured using a microplate reader (Bio-Rad, Hercules, USA) at 450 nm.

### 2.4 Western Blotting

Next, western blotting was performed to investigate the changes in N2a/APP cells after gastrodin treatment in terms of the level of amyloid precursor protein-derived C-terminal fragments (APP-CTFs), presynaptic proteins (SNAP25 and synaptophysin) and postsynaptic proteins (PSD95); phosphorylated-tau (p-tau) and total-tau (t-tau); phosphorylated-ERK (p-ERK) and total-ERK (t-ERK); phosphorylated-JNK (p-JNK) and total-JNK (t-JNK); phosphorylated-GSK3 b (p-GSK3b) and total-GSK3b (t-GSK3b) proteins as described previously [56]. The antibodies used are listed in **Supplementary Table 1**. Equal amounts of protein per lane (20–50 μg) were loaded onto 4%–12% sodium dodecyl sulfate–polyacrylamide gels (Absin, China) for 40 min using an electrophoresis apparatus (Bio-Rad, USA) and transferred to polyvinylidene difluoride membranes (Millipore, USA) for 7 min. The membranes were incubated in 5% skim milk powder (weight/volume) for 2 hours at room temperature, followed by overnight incubation at 4 °C with primary antibody. Subsequently, the membranes were washed three times with TBST (20% Tween-20 added in Tris buffered saline) and incubated with secondary antibody for 1 hour. Anti-β-tubulin or anti-glyceraldehyde 3-phosphate dehydrogenase (GAPDH) antibodies were used as loading controls. Each experiment was repeated at least 3 times. An imaging system (SYNGENE, UK) was used to evaluate the protein expression of each lane, and the band density was analysed by ImageJ software (ImageJ, National Institute of Mental Health, USA).

### 2.5 Flow cytometry

Flow cytometry using dichlorodihydrofluorescein diacetate (DCFH-DA) was used to detect the level of ROS in N2a/APP cells, as described previously [19]. Briefly, N2a/APP cells were treated with gastrodin at different concentrations (1 μM, 5 μM, and 10 μM) (n = 3) for 24 h and incubated with 10 μM DCFH-DA for 30 min at 37°C. Cells were rotated gently, washed and resuspended in phosphate-buffered saline (PBS, pH 7.4). The mean fluorescence intensity was detected by a FACS VerseTM flow cytometer (BD Biosciences, USA). Data were analysed by FlowJo software (FlowJo X 10.0.7r2, BD Biosciences, USA).

Flow cytometry using the fluorescent dye JC-1, 5,5′,6,6′-tetrachloro-1,1′,3,3′-tetraethylbenzimidazolylcarbocyanine iodide, was used to measure changes in the mitochondrial membrane potential (ΔΨm), as described previously [19]. N2a/APP cells (1 × 10^6^) were seeded in 6-well plates and treated with 1 μM, 5 μM, and 10 μM gastrodin (n = 3). Then, the cells were harvested, washed with PBS and incubated with fresh medium containing JC-1 at 37°C for 20 min. The cells were then washed with staining buffer twice and detected using a FACS Verse TM flow cytometer. The monomer/aggregate ratio (green fluorescence/red fluorescence ratio) was analysed by FACSuite software (BD Biosciences, USA).

### 2.7 Immunofluorescence Staining

Immunofluorescence staining of 4-hydroxynonenal (4-HNE) for the quantification of the level of lipid peroxidation and 8-hydroxydeoxyguanosine (8-OHdG) for the level of DNA oxidation was performed as described previously [57]. N2a/APP cells (1×10^4^) seeded on 14-16 mm coverslips were exposed to different concentrations of gastrodin for 24 h. After that, the cells were fixed with 4% paraformaldehyde for 20 minutes, permeabilized with 0.3% Triton X-100 for 5 minutes, and blocked with 5% goat serum for 60 minutes at room temperature. Next, the cells were sequentially incubated with primary antibodies against 4-HNE (1:150) or 8-OHdG (1:200) at 4°C overnight and then with appropriate secondary antibodies conjugated with Alexa Fluor 488 (1:200) or Alexa Fluor 549 (1:200) for 1 hour at 37°C. PBS was used for washes. Cells were mounted with a medium containing anti◻fading agents and 4′,6◻diamidino◻2◻phenylindole (DAPI) for counterstaining. Confocal images of immunofluorescence-stained cells were obtained using a 100× oil objective on a confocal microscope (Olympus, Japan). Confocal images were processed with OlyVIA software (OLYMPUS OlyVIA3.3, Olympus, Japan) and analysed by using ImageJ (NIH, United States).

### 2.8 Statistical Analysis

Statistical results are expressed as mean ± standard error (SEM) (n = 3-6). One-way ANOVA and Tukey’s post hoc analysis were used for multiple-group comparisons using GraphPad Prism (Version 9.0, GraphPad Software, LLC, USA). A p value <0.05 was considered statistically significant.

## 3. Results

### 3.1 Gastrodin ameliorated synaptic impairment in N2a/APP cells

First, we determined the optimal dosage for gastrodin treatment and evaluated the effects of gastrodin on cell viability by treating N2a/APP cells with gastrodin at concentrations ranging from 0.2 μM to 10 μM. No loss of cell viability was observed in N2a/APP cells treated with 0.2 μM to 5 μM gastrodin. Treatment with 10 μM gastrodin resulted in a 15% reduction in cell viability (p = 0.0007, vs. control) (**Fig. 1a**).

**Figure 1.**
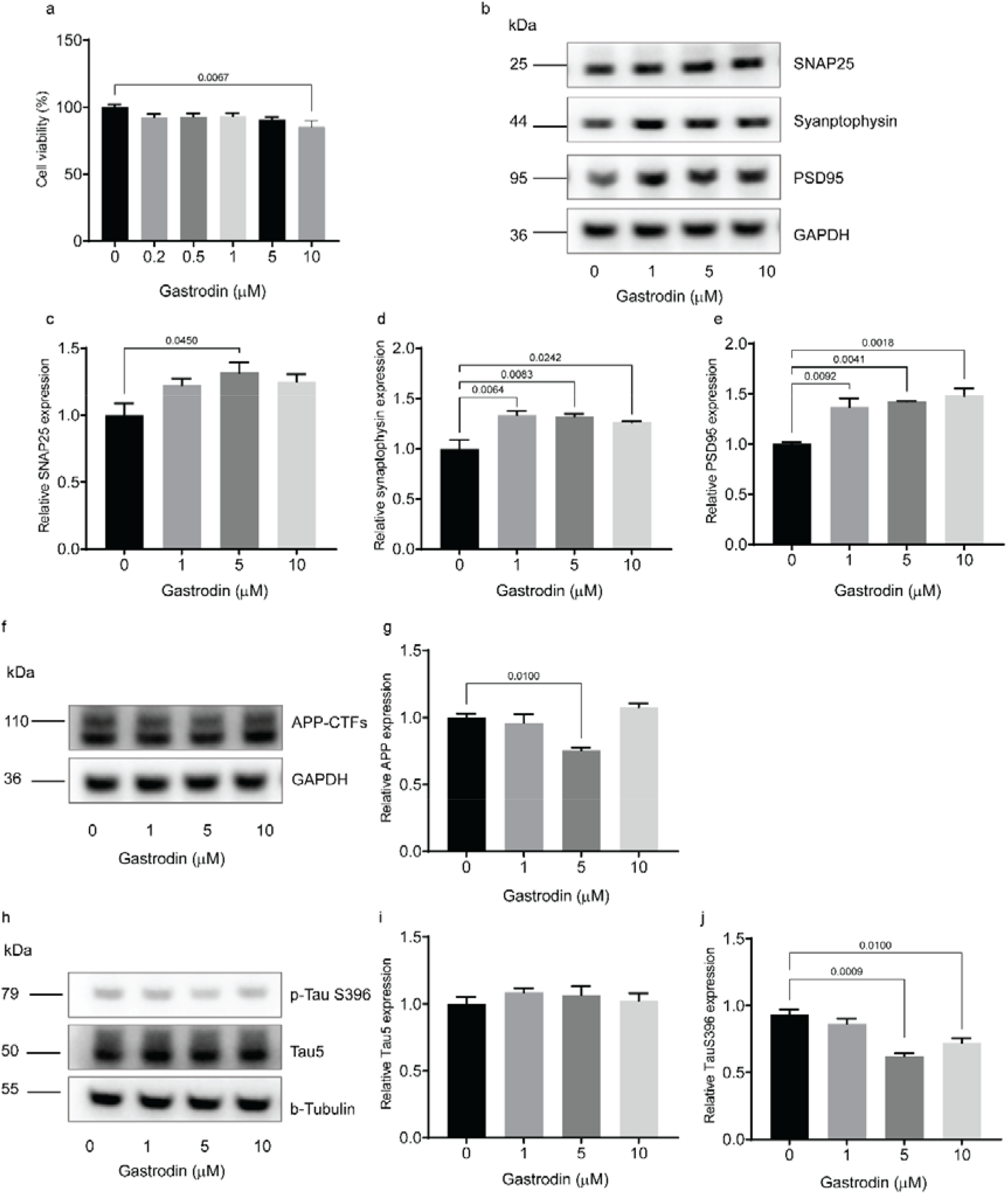
Gastrodin increased synaptic protein expression, reduced the levels of APP-CTFs and phospho-Tau S396 in in N2A/APP cells. (**a, b, d**) Cell viability test for gastrodin at different concentrations assessed by using the CCK-8 assay (n =6). (**b-e**) Representative blots and quantification of the expression of presynaptic SNAP25, synaptophysin, and postsynaptic PSD95 in N2A/APP cells after gastrodin treatment (n = 3). (**f**, **g**) Representative blots and quantification of the expression of APP-CTFs in N2A/APP cells after gastrodin treatment (n = 3). (**h-j**) Representative blots and quantification of the expression of total tau (Tau5) and phosphorylated TauS396 in N2A/APP cells after gastrodin treatment (n = 3). Values were normalized to the control. Data are presented as the mean ± SEM. One-way ANOVA, Tukey’multiple comparison

Next, we investigated the potential effects of gastrodin on synaptic plasticity-related proteins, including presynaptic terminal proteins (synaptophysin and SNAP25) and postsynaptic density protein (PSD95), by using western blotting. We found that treatment with 5 μM gastrodin (but not 1 or 10 μM) significantly increased the expression levels of presynaptic SNAP25 by 32% in N2a/APP cells (p = 0.045, vs. control) (**Figs. 1b.c**). Treatment with gastrodin elevated the levels of synaptophysin by approx. 30% (1 μM p = 0.0064, 5 μM p = 0.0083, 10 μM p = 0.0242, vs. control) and the levels of postsynaptic PSD95 by approx. 40% (1 μM p = 0.0092, 5 μM p = 0.0041, 10 μM p = 0.0018, vs. control) (**Figs. 1b, d, e**).

### 3.3 Gastrodin decreased the expression of APP-CTFs and phosphorylated tau protein in N2a/APP cells

In the amyloidogenic pathway, APP is sequentially cleaved by β-secretases and produces soluble peptide APPβ and APP-C-terminal fragments (CTFs). Recent studies have suggested an association between APP-CTF levels and mitochondrial dysfunction and neurotoxicity in models of AD [58]. Here, we assessed the influence of gastrodin treatment on the expression of APP-CTFs in N2a/APP cells by using western blotting. Treatment with 5 μM gastrodin decreased the expression of APP-CTFs by 25% in N2a/APP cells (p = 0.01, vs. control), and no changes were observed with treatment using 1 or 10 μM gastrodin (**Figs. 1f, g**).

Furthermore, we investigated the effect of gastrodin treatment on tau and tau phosphorylation based on the modulatory effect reported in previous studies [35, 59]. We found that treatment with gastrodin decreased the expression of phosphorylated tau protein at Ser396 (5 μM p = 0.0009, 10 μM p = 0.0045, vs. control). There was no change in the total tau level with treatment with different concentrations of gastrodin. (**Figs. 1h-j**).

### 3.4 Gastrodin inhibited the phosphorylation of GSK-3β and activated the phosphorylation of ERK1/2 and JNK in N2a/APP cells

We hypothesized that the ERK1/2, GSK-3β, and JNK signaling pathways were involved in the neuroprotective effects of gastrodin. We assessed the influence of gastrodin pretreatment on p-ERK1/2, t-ERK1/2, p-GSK-3β, t-GSK-3β, t-JNK, and p-JNK in N2a/APP cells by using western blotting. We found that gastrodin significantly decreased the level of phosphorylated GSK-3β in N2a/APP cells at 5 μM (p = 0.0297, vs. control) and 10 μM (p = 0.0017, vs. control) (**Figs. 2a, b**). No alteration in the level of t-GSK-3β was detected with gastrodin treatment (**Fig. 2c**). Gastrodin significantly increased the level of phosphorylated ERK1/2 in N2a/APP cells at 5 μM (p = 0.0067, vs. control) (**Figs. 2d, e**). No alteration in the level of t-ERK1/2 was detected with gastrodin treatment (**Fig. 2f**). Gastrodin significantly increased the level of phosphorylated JNK in N2a/APP cells at 5 μM (p = 0.0206, vs. control) (**Figs. 2g, h**). No alteration in the level of t-ERK1/2 was detected with gastrodin treatment (**Figs. 2i**).

**Figure 2.**
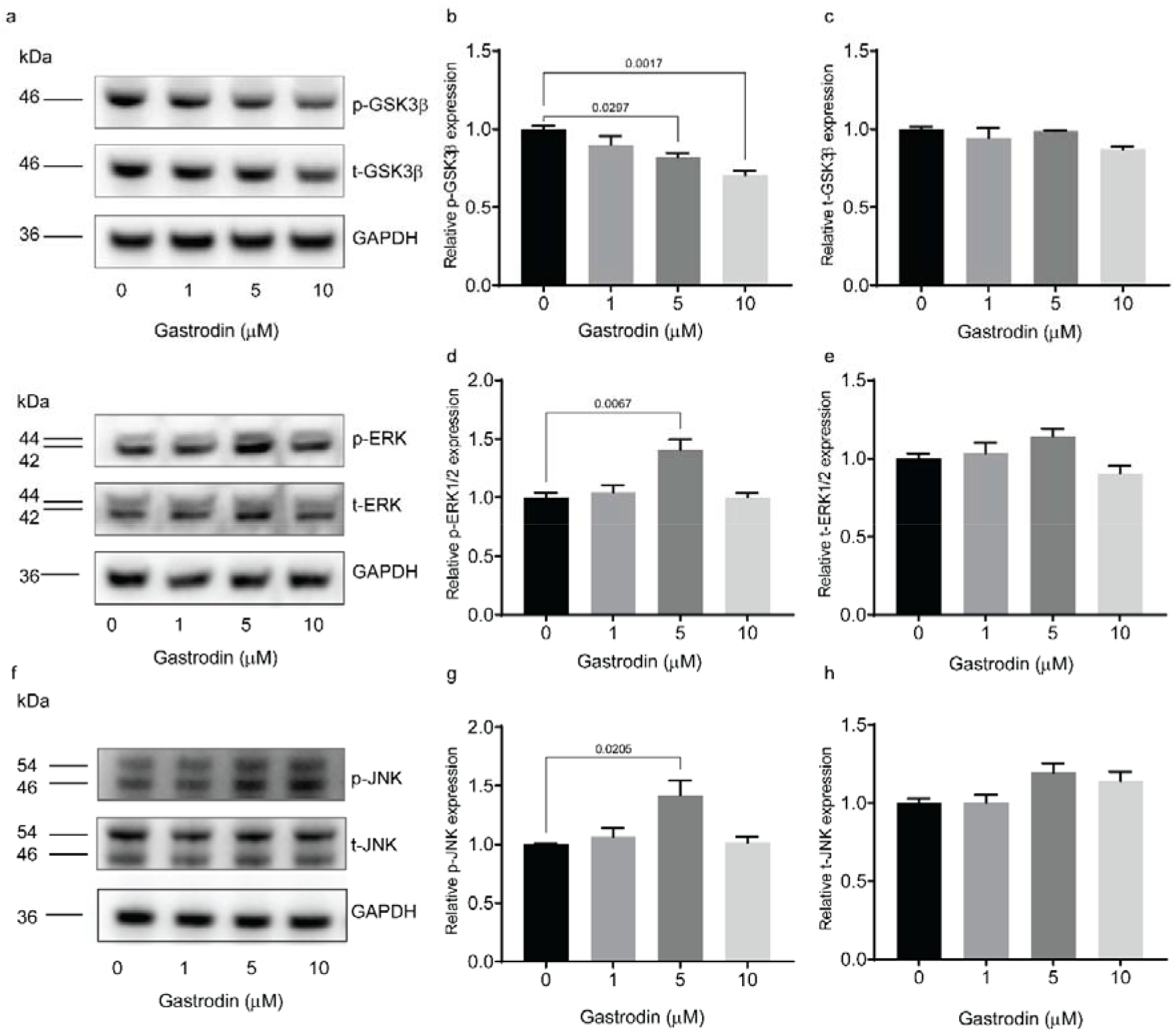
Gastrodin inhibited the hyperactivation of phosphorylated GSK-3β and enhanced the phosphorylation of ERK1/2 and JNK in N2a/APP cells. (**a-c**) Representative blots and quantification of the level of p-GSK-3β and t-GSK-3β in N2a/APP cells after gastrodin treatment.(**e-g**) Representative blots and quantification of the level of p-ERK1/2 and t-ERK1/2 in N2a/APP cells after gastrodin treatment (n = 3). (**h-j**) Representative blots and quantification of the level of p-JNK and t-JNK in N2a/APP cells. after gastrodin treatment(n=3). Values were normalized to the control. Data are presented as the mean ± SEM, one-way ANOVA, Tukey’s multiple comparison.

### 3.5 Gastrodin attenuated the production of ROS and reestablished mitochondrial membrane potential (ΔΨm) in N2a/APP cells

Next, we examined the effect of gastrodin on the production of ROS in N2a/APP cells by flow cytometry using the fluorescence probe DCFH-DA. Gastrodin treatment at 5 μM (p = 0.0001, vs. control) and 10 μM (p = 0.0485, vs. control) decreased the ROS level in N2a/APP cells (**Figs. 3a-d, i**). In addition, we assessed the effect of gastrodin on the mitochondrial membrane potential (ΔΨm) in N2a/APP cells by flow cytometry using JC-1 staining. JC-1 aggregates in coupled mitochondria emit red fluorescence, while the JC-1 monomeric form emits green fluorescence when cells lose ΔΨm. Gastrodin treatment decreased the JC-1 monomer/aggregate ratio (green/red fluorescence intensity ratio) at 1 μM (p = 0.0643, vs. control), 5 μM (p = 0.0214, vs. control) and 10 μM (p = 0.001, vs. control), implying the reestablishment of ΔΨm in N2a/APP cells (**Figs. 3e-h, j**).

**Figure 3.**
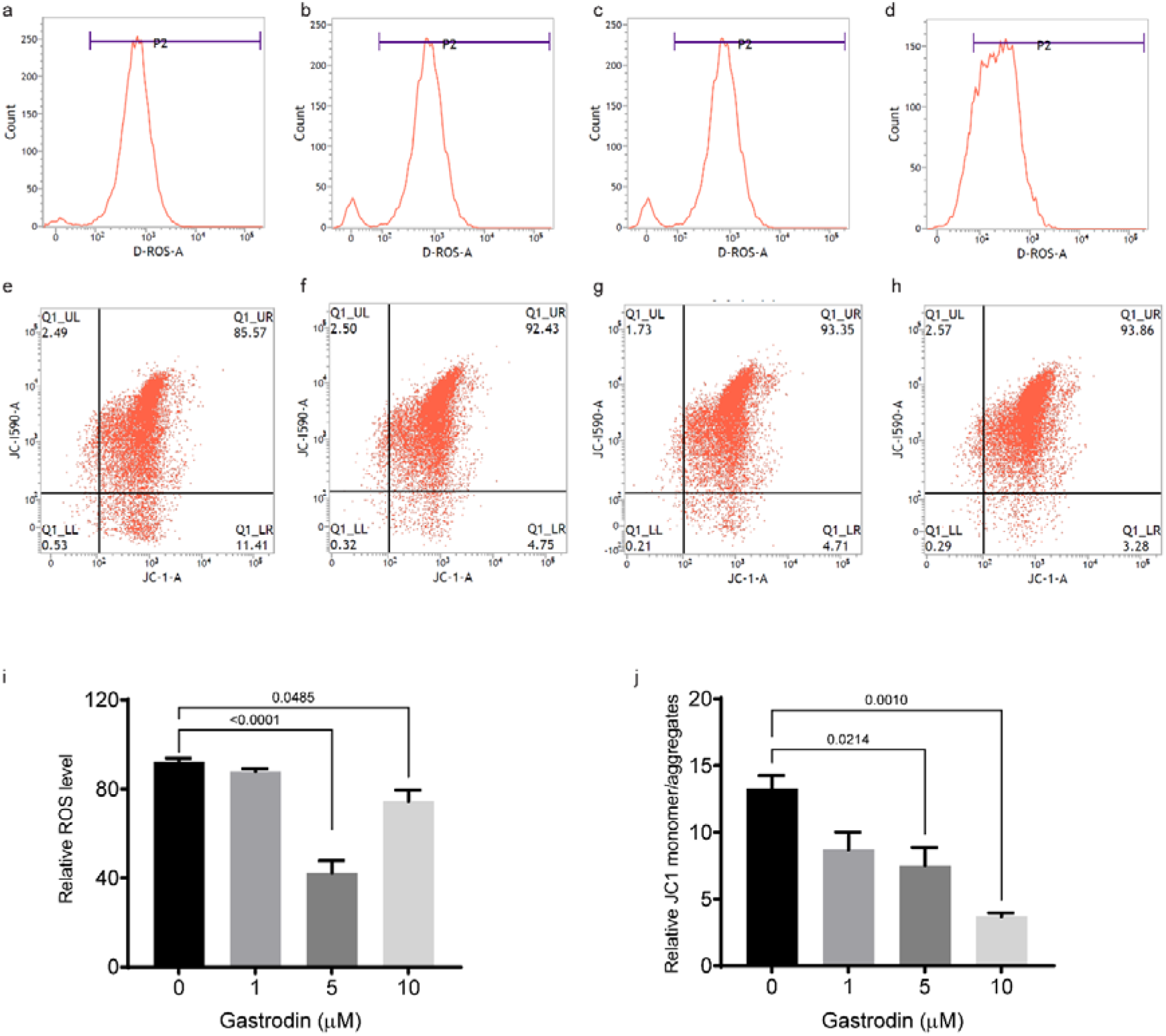
Gastrodin attenuated the production of ROS and reestablished mitochondrial membrane potential (ΔΨm) in N2a/APP cells. (**a-d**, **i**) Analysis of ROS with DCFH-DA by flow cytometry, (a) Control, (b) 1 μM gastrodin treatment, (c) 5 μM gastrodin treatment, (d) 10 μM gastrodin treatment, (i) Quantification of ROS in N2a/APP cells after gastrodin treatment (n = 3). (**e-h**, **j**) Analysis of ΔΨm with JC-1 staining by flow cytometry (n=3). (e) Control, (f) 1 μM gastrodin treatment, (g) 5 μM gastrodin treatment, (h) 10 μM gastrodin treatment, (j) Quantification of monomer/aggregates ratio indicated by green/red fluorescence intensity in N2a/APP cells after gastrodin treatment (n = 3). Data are presented as the mean ± SEM, one-way ANOVA, Tukey’s multiple comparison.

### 3.6 Gastrodin ameliorated oxidative stress in N2a/APP cells

Next, we evaluated the effects of gastrodin on oxidative stress, including lipid peroxidation (by using 4-HNE staining) and DNA damage (by using 8-OHdG staining), in N2a/APP cells. Treatment with gastrodin (1. 5, 10 μM) reduced the 4-HNE fluorescence intensity produced by lipid peroxidation in N2a/APP cells in a dose◻dependent manner (p<0.0001 for all groups, vs.) (**Figs. 4a, b**). We found that treatment with gastrodin at 5 and 10 μM (but not 1 μM) significantly reduced the fluorescence intensity of 8-OHdG in N2a/APP cells by approximately 64% (p < 0.0001, vs. control) and 87% (p < 0.0001, vs. control), respectively (**Figs. 4c, d**).

**Figure 4.**
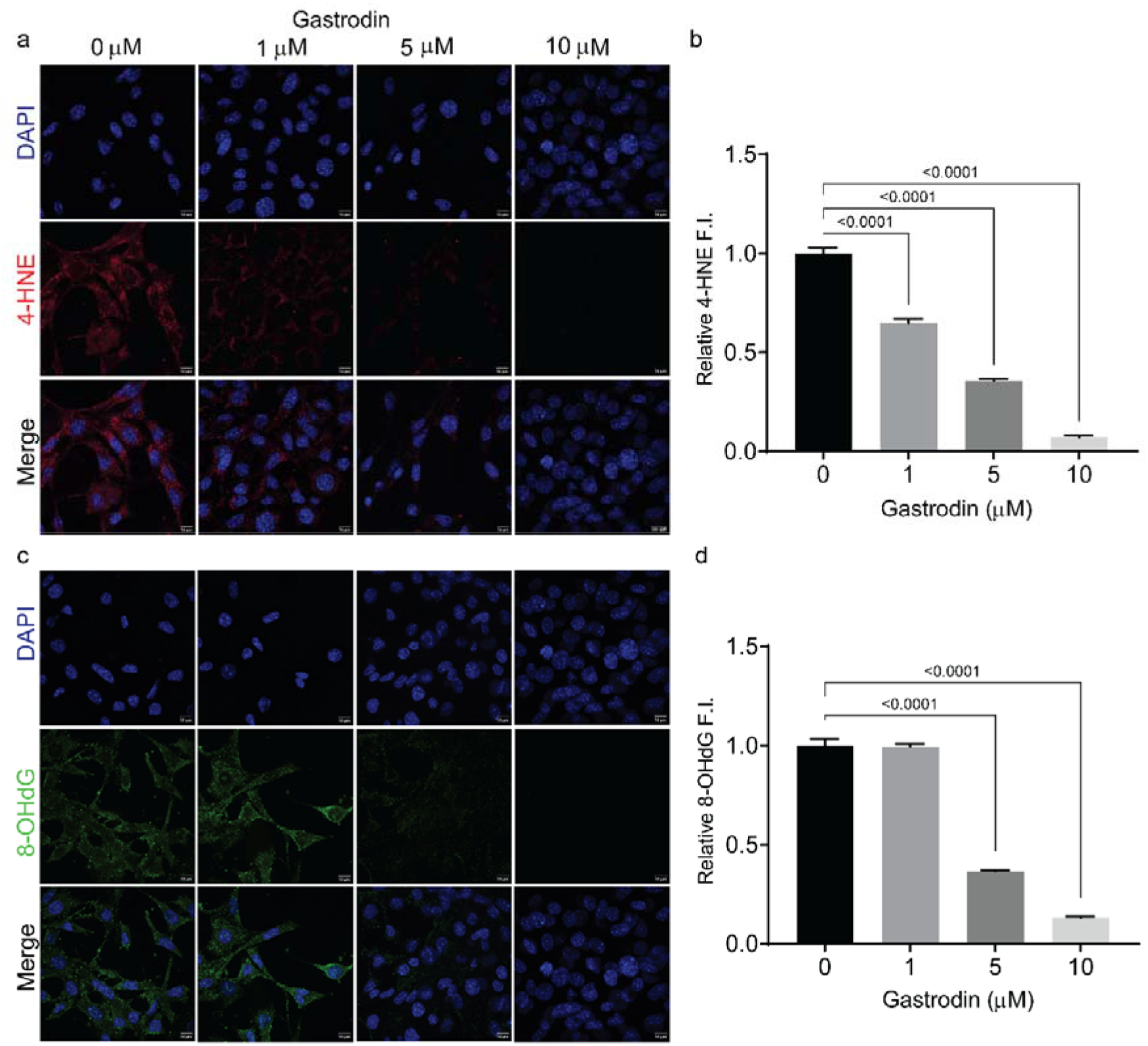
Gastrodin suppresses lipid peroxidation and DNA oxidation in N2a/APP cells. (**a**, **b**) Representative confocal images and quantification of the immunoreactivity of 4-HNE in N2a/APP cells with or without gastrodin treatment (n = 3). Nuclei are counterstained with DAPI (blue). (**c**, **d**) Representative confocal images and quantification of the immunoreactivity of 8-OHdG (n = 3). Nuclei are counterstained with DAPI. Scale bar = 10 μm (a-d), 50 cells were analysed in each group. One-way ANOVA, Tukey’s multiple comparison.

## 4. Discussion

Here, we demonstrated that gastrodin attenuated APP-CTFs levels, p-TauS396, synaptic dysfunction, ROS generation, and oxidative stress and restored mitochondrial function. Furthermore, we found that gastrodin treatment led to increases in p-ERK and p-JNK expression and a reduction in p-GSK-3β expression in N2a/APP cell model of AD.

The effect of gastrodin on APP processing has been reported to suppress BACE1 expression [35, 37] and promote neuroprotective α-secretase [60]. Here, we demonstrated that gastrodin treatment reduced the level of APP-CTFs expression in N2a/APP cells. Recent studies have shown that the accumulation of APP-CTFs is associated with lysosome dysfunction [61], synaptic impairment, and mitochondrial and mitophagy dysfunction in AD-induced neural stem cells [58], cultured rat basal forebrain cholinergic neurons [62], 3×Tg AD mice and brains from patients with AD [63]. Impaired synaptic function and long-term memory were reported in mice overexpressing C99 fragments or injected with APP-CTFs showed [7, 8]. Higher levels of APP-CTFs were detected in the cerebrospinal fluid of autosomal AD patients carrying PSEN1 mutations [64].

Moreover, we showed that gastrodin treatment reversed the decreased expression levels of the synaptic proteins SNAP 25, synaptophysin, and PSD-95 in both N2a/APP cells. The effect of gastrodin on synaptic proteins has not much reported. In an Aβ injected mouse model of AD, gastrodin treatment led to an increasing trend in the levels of BDNF, PSD-95, and synaptophysin [65]. A previous study demonstrated that gastrodin treatment increased the levels of brain-derived neurotrophic factor (BDNF), PSD-95, and synaptophysin in a mouse model of Parkinson’s disease [66].

Tau phosphorylation plays an important role regulating tau function, and involved in the regulation of a range of signaling pathways, including GSK-3β [67]. We showed that gastrodin treatment could reduce the level of tau phosphorylated at Ser396, while the total tau level remained stable. Our finding is in line with a previous report that gastrodin treatment reduced tau phosphorylation at Ser199, Thr231, and Ser396 in APP/PS1 transgenic mice of AD [35]. The effect of gastrodin on p-tau has been previously shown in aged mice [26], lead-induced brain injury mice [50], a Drosophila PINK1 model of Parkinson’s disease [68], and rats OF vascular dementia [59].

Reducing oxidative stress is a therapeutic target for slowing neurodegeneration in AD [69]. We observed that gastrodin treatment reduced the overproduction of ROS, reestablished the mitochondrial membrane potential, and attenuated lipid peroxidation (4-HNE) and DNA damage (8-OHdG) in N2a/APP cells. Increased ROS and oxidative stress play an important role in AD and have been reported in the AD cellular model [70, 71], animal models and brains from patients with AD [69, 72]. In addition ROS-induced oxidative stress causes mitochondrial dysfunction and contributes to Aβ production and aggregation [73] and tau phosphorylation [20], thus facilitating AD pathogenesis [23, 72, 74, 75]. The effect of gastrodin on mitochondrial function, associated redox and bioenergetics effect, has been demonstrated in Aβ-injected mouse models as well as in the SH-SY5Y cells [76, 77]. Further analysis of Mitochondria function such as Mitochondrial fusion and fission protein level will be of interest.

We found that gastrodin treatment increased the levels of p-ERK and p-JNK and reduced the level of p-GSK-3β, while t-ERK, t-GSK-3β and t-JNK remained stable compared to the control group. Our results are in line with earlier findings of the gastrodin effect mediated via downregulation of GSK-3β phosphorylation [26, 35, 48, 78, 79] and upregulation of ERK1/2 phosphorylation [34, 46, 47, 78, 80, 81] in AD cellular and animal models. Intraneuronal Aβ_1–42_ accumulation has been shown to lead to a reduction in the phosphorylation of ERK 1/2 [82]. Earlier studies have shown the involvement of the GSK-3β/ERK1/2 pathway in the antioxidant properties of gastrodin [83]. GSK-3β is a vital regulator involved in neuronal processes, including axonal transport and synaptic function [84, 85]. Overexpression and phosphorylation of GSK-3β is detected in the brains of AD patients [53, 54] and promotes tau hyperphosphorylation and cognitive deficits in animal models of AD [86–89] Reducing abnormal GSK-3β expression reversed memory deficits and decreased neuronal death and tau hyperphosphorylation in a mouse model [90]. Similar to our finding, a previous study in APP/PS1 mice showed that gastrodin treatment decreased the level of p-GSK3β (including Ser396 sites) [35]. The JNK pathway is critical in cell death and apoptosis and has been less investigated in regard to AD pathology. An earlier study in the brain and cerebral spinal fluid of patients with AD showed that increased levels of phosphorylated JNK3 correlate with the progression of AD [91]. A previous study demonstrated that inhibition of JNK prevented Aβ oligomer-induced synapse loss and neuronal cell death in TgCRND8 mice with AD amyloidosis [92]. To date, only one study has investigated the involvement of JNK phosphorylation by gastrodin in response to Aβ42 in neural progenitor cells [47], in which a downregulation effect was reported. The potential reason for the difference with our results could be the different duration dosages of gastrodin treatment and the cell model utilized. Further analysis of JNK subtypes is needed to elucidate the involvement of this pathway.

There are several limitations in our study. 1) We have only examined the involvement of ERK, GSK3β, JNK pathways in the effect of gastrodin. Several other pathways mediating the effect of gastrodin might be also involved such as inhibition of caspase-3 Aβ-induced toxicity, suppression of intracellular calcium elevation [93], PKR/eIF2a [37], TLR4/NF-κB [36] and MAPK/PI3K [49], PDE9-cGMP-PKG [94], and NLRP3 inflammasome activation [95]. 2) Treatment using gastrodin at 5 μM was effective dosage in the different assessment with no cytotoxicity. However dosage response pattern in the treatment effect was not fully clear. Further optimization in the dosage is needed.

## 5. Conclusion

In conclusion, we show that gastrodin exerts neuroprotective effects in N2a/APP cells by attenuating p-GSK-3β, increasing p-ERK1/2 and p-JNK expression, attenuating APP-CTFs and tau levels, and alleviating mitochondrial dysfunction and oxidative stress in the APP/PS1 cell model of AD.

## Author Contributions

ZT, RN, and XLQ contributed to the conception and study design. YQP, MG and TZ performed the experiments. ZT, YQP and YX contributed to data collection and data analysis. RN and ZT interpreted the data. XLQ, YQP, RN, ZYC and ZT wrote the manuscript. All authors approved the manuscript before submission.

## References

1. Scheltens P, De Strooper B, Kivipelto M, Holstege H, Chételat G, Teunissen CE, Cummings J, van der Flier WM: Alzheimer’s disease. The Lancet 2021, 397(10284):1577–1590.

2. Haass C, Willem M: Secreted APP Modulates Synaptic Activity: A Novel Target for Therapeutic Intervention? Neuron 2019, 101(4):557–559.

3. Selkoe DJ, Hardy J: The amyloid hypothesis of Alzheimer’s disease at 25 years. EMBO Mol Med 2016, 8(6):595–608.

4. Multhaup G, Huber O, Buee L, Galas MC: Amyloid Precursor Protein (APP) Metabolites APP Intracellular Fragment (AICD), Abeta42, and Tau in Nuclear Roles. The Journal of biological chemistry 2015, 290(39):23515–23522.

5. Butterfield DA, Boyd-Kimball D: Oxidative Stress, Amyloid-β Peptide, and Altered Key Molecular Pathways in the Pathogenesis and Progression of Alzheimer’s Disease. Journal of Alzheimer’s disease: JAD 2018, 62(3):1345–1367.

6. Butterfield DA, Swomley AM, Sultana R: Amyloid beta-Peptide (1-42)-Induced Oxidative Stress in Alzheimer Disease: Importance in Disease Pathogenesis and Progression. Antioxidants & redox signaling 2013, 19(8):823–835.

7. Cheignon C, Tomas M, Bonnefont-Rousselot D, Faller P, Hureau C, Collin F: Oxidative stress and the amyloid beta peptide in Alzheimer’s disease. Redox biology 2018, 14:450–464.

8. Busche MA, Hyman BT: Synergy between amyloid-β and tau in Alzheimer’s disease. Nat Neurosci 2020, 23(10):1183–1193.

9. Denechaud M, Geurs S, Comptdaer T, Bégard S, Garcia-Núñez A, Pechereau LA, Bouillet T, Vermeiren Y, De Deyn PP, Perbet R et al: Tau promotes oxidative stress-associated cycling neurons in S phase as a pro-survival mechanism: Possible implication for Alzheimer’s disease. Prog Neurobiol 2022:102386.

10. Kandimalla R, Manczak M, Yin X, Wang R, Reddy PH: Hippocampal phosphorylated tau induced cognitive decline, dendritic spine loss and mitochondrial abnormalities in a mouse model of Alzheimer’s disease. Human molecular genetics 2018, 27(1):30–40.

11. Wang W, Zhao F, Ma X, Perry G, Zhu X: Mitochondria dysfunction in the pathogenesis of Alzheimer’s disease: recent advances. Mol Neurodegener 2020, 15(1):30.

12. Wang X, Su B, Siedlak SL, Moreira PI, Fujioka H, Wang Y, Casadesus G, Zhu X: Amyloid-beta overproduction causes abnormal mitochondrial dynamics via differential modulation of mitochondrial fission/fusion proteins. Proc Natl Acad Sci U S A 2008, 105(49):19318–19323.

13. Islam MT: Oxidative stress and mitochondrial dysfunction-linked neurodegenerative disorders. Neurological research 2017, 39(1):73–82.

14. Reddy VP, Zhu X, Perry G, Smith MA: Oxidative stress in diabetes and Alzheimer’s disease. Journal of Alzheimer’s disease: JAD 2009, 16(4):763–774.

15. Zhang J, Liu L, Zhang Y, Yuan Y, Miao Z, Lu K, Zhang X, Ni R, Zhang H, Zhao Y et al: ChemR23 signaling ameliorates cognitive impairments in diabetic mice via dampening oxidative stress and NLRP3 inflammasome activation. Redox Biology 2022, 58:102554.

16. Ionescu-Tucker A, Cotman CW: Emerging roles of oxidative stress in brain aging and Alzheimer’s disease. Neurobiology of aging 2021, 107:86–95.

17. Kim GH, Kim JE, Rhie SJ, Yoon S: The Role of Oxidative Stress in Neurodegenerative Diseases. Experimental neurobiology 2015, 24(4):325–340.

18. Cieślik M, Czapski GA, Wójtowicz S, Wieczorek I, Wencel PL, Strosznajder RP, Jaber V, Lukiw WJ, Strosznajder JB: Alterations of Transcription of Genes Coding Anti-oxidative and Mitochondria-Related Proteins in Amyloid β Toxicity: Relevance to Alzheimer’s Disease. Mol Neurobiol 2020, 57(3):1374–1388.

19. Lai C, Chen Z, Ding Y, Chen Q, Su S, Liu H, Ni R, Tang Z: Rapamycin Attenuated Zinc-Induced Tau Phosphorylation and Oxidative Stress in Rats: Involvement of Dual mTOR/p70S6K and Nrf2/HO-1 Pathways. Frontiers in immunology 2022, 13:782434.

20. Alavi Naini SM, Soussi-Yanicostas N: Tau Hyperphosphorylation and Oxidative Stress, a Critical Vicious Circle in Neurodegenerative Tauopathies? Oxid Med Cell Longev 2015, 2015:151979.

21. Liu Z, Li T, Li P, Wei N, Zhao Z, Liang H, Ji X, Chen W, Xue M, Wei J: The Ambiguous Relationship of Oxidative Stress, Tau Hyperphosphorylation, and Autophagy Dysfunction in Alzheimer’s Disease. Oxidative medicine and cellular longevity 2015, 2015:352723.

22. Haddadi M, Jahromi SR, Sagar BK, Patil RK, Shivanandappa T, Ramesh SR: Brain aging, memory impairment and oxidative stress: a study in Drosophila melanogaster. Behavioural brain research 2014, 259:60–69.

23. Meraz-Ríos MA, Franco-Bocanegra D, Toral Rios D, Campos-Peña V: Early onset Alzheimer’s disease and oxidative stress. Oxidative medicine and cellular longevity 2014, 2014:375968.

24. Liu Y, Gao J, Peng M, Meng H, Ma H, Cai P, Xu Y, Zhao Q, Si G: A Review on Central Nervous System Effects of Gastrodin. Frontiers in pharmacology 2018, 9:24.

25. Hu Y, Li C, Shen W: Gastrodin alleviates memory deficits and reduces neuropathology in a mouse model of Alzheimer’s disease. Neuropathology: official journal of the Japanese Society of Neuropathology 2014, 34(4):370–377.

26. Wang X, Chen L, Xu Y, Wang W, Wang Y, Zhang Z, Zheng J, Bao H: Gastrodin alleviates perioperative neurocognitive dysfunction of aged mice by suppressing neuroinflammation. European journal of pharmacology 2021, 892:173734.

27. Guo J, Zhang XL, Bao ZR, Yang XK, Li LS, Zi Y, Li F, Wu CY, Li JJ, Yuan Y: Gastrodin Regulates the Notch Signaling Pathway and Sirt3 in Activated Microglia in Cerebral Hypoxic-Ischemia Neonatal Rats and in Activated BV-2 Microglia. Neuromolecular medicine 2021, 23(3):348–362.

28. Yuan B, Huang H, Qu S, Zhang H, Lin J, Jin L, Yang S, Zeng Z: Gastrodin Pretreatment Protects Liver Against Ischemia-Reperfusion Injury via Activation of the Nrf2/HO-1 Pathway. The American journal of Chinese medicine 2020, 48(5):1159–1178.

29. Zhang F, Deng CK, Huang YJ, Miao YH, Wang YY, Zhang Y, Qian ZY, Zhang WQ, Zhou RD, Lei B et al: Early Intervention of Gastrodin Improved Motor Learning in Diabetic Rats Through Ameliorating Vascular Dysfunction. Neurochemical research 2020, 45(8):1769–1780.

30. Li Q, Li L, Yu M, Zheng M, Li Y, Yang J, Dai M, Zhong L, Sun L, Lu D: Elastomeric polyurethane porous film functionalized with gastrodin for peripheral nerve regeneration. J Biomed Mater Res A 2020, 108(8):1713–1725.

31. Peng Z, Wang H, Zhang R, Chen Y, Xue F, Nie H, Chen Y, Wu D, Wang Y, Wang H et al: Gastrodin ameliorates anxiety-like behaviors and inhibits IL-1beta level and p38 MAPK phosphorylation of hippocampus in the rat model of posttraumatic stress disorder. Physiol Res 2013, 62(5):537–545.

32. Li Y, Zhang Z: Gastrodin improves cognitive dysfunction and decreases oxidative stress in vascular dementia rats induced by chronic ischemia. International journal of clinical and experimental pathology 2015, 8(11):14099–14109.

33. Wang X, Yan S, Wang A, Li Y, Zhang F: Gastrodin ameliorates memory deficits in 3,3’-iminodipropionitrile-induced rats: possible involvement of dopaminergic system. Neurochemical research 2014, 39(8):1458–1466.

34. Wang XL, Xing GH, Hong B, Li XM, Zou Y, Zhang XJ, Dong MX: Gastrodin prevents motor deficits and oxidative stress in the MPTP mouse model of Parkinson’s disease: involvement of ERK1/2-Nrf2 signaling pathway. Life sciences 2014, 114(2):77–85.

35. Zeng Y-Q, Gu J-H, Chen L, Zhang T-T, Zhou X-F: Gastrodin as a multi-target protective compound reverses learning memory deficits and AD-like pathology in APP/PS1 transgenic mice. Journal of Functional Foods 2021, 77:104324.

36. Fasina OB, Wang J, Mo J, Osada H, Ohno H, Pan W, Xiang L, Qi J: Gastrodin From Gastrodia elata Enhances Cognitive Function and Neuroprotection of AD Mice via the Regulation of Gut Microbiota Composition and Inhibition of Neuron Inflammation. Front Pharmacol 2022, 13:814271.

37. Zhang JS, Zhou SF, Wang Q, Guo JN, Liang HM, Deng JB, He WY: Gastrodin suppresses BACE1 expression under oxidative stress condition via inhibition of the PKR/eIF2α pathway in Alzheimer’s disease. Neuroscience 2016, 325:1–9.

38. Hao S, Yang Y, Han A, Chen J, Luo X, Fang G, Liu J, Wang S: Glycosides and Their Corresponding Small Molecules Inhibit Aggregation and Alleviate Cytotoxicity of Aβ40. ACS Chem Neurosci 2022, 13(6):766–775.

39. Luo K, Wang Y, Chen WS, Feng X, Liao Y, Chen S, Liu Y, Liao C, Chen M, Ao L: Treatment Combining Focused Ultrasound with Gastrodin Alleviates Memory Deficit and Neuropathology in an Alzheimer’s Disease-Like Experimental Mouse Model. Neural Plast 2022, 2022:5241449.

40. Liu XC, Wu CZ, Hu XF, Wang TL, Jin XP, Ke SF, Wang E, Wu G: Gastrodin Attenuates Neuronal Apoptosis and Neurological Deficits after Experimental Intracerebral Hemorrhage. J Stroke Cerebrovasc Dis 2020, 29(1):104483.

41. Liang WZ, Jan CR, Hsu SS: Cytotoxic effects of gastrodin extracted from the rhizome of Gastrodia elata Blume in glioblastoma cells, but not in normal astrocytes, via the induction of oxidative stress-associated apoptosis that involved cell cycle arrest and p53 activation. Food and chemical toxicology: an international journal published for the British Industrial Biological Research Association 2017, 107(Pt A):280–292.

42. Cheng QQ, Wan YW, Yang WM, Tian MH, Wang YC, He HY, Zhang WD, Liu X: Gastrodin protects H9c2 cardiomyocytes against oxidative injury by ameliorating imbalanced mitochondrial dynamics and mitochondrial dysfunction. Acta pharmacologica Sinica 2020, 41(10):1314–1327.

43. Sadeghian Z, Eyvari-Brooshghalan S, Sabahi M, Nourouzi N, Haddadi R: Post treatment with Gastrodin suppresses oxidative stress and attenuates motor disorders following 6-OHDA induced Parkinson disease. Neuroscience letters 2022, 790:136884.

44. Huang L, Shao M, Zhu Y: Gastrodin inhibits high glucose-induced inflammation, oxidative stress and apoptosis in podocytes by activating the AMPK/Nrf2 signaling pathway. Experimental and therapeutic medicine 2022, 23(2):168.

45. Jiang G, Hu Y, Liu L, Cai J, Peng C, Li Q: Gastrodin protects against MPP(+)-induced oxidative stress by up regulates heme oxygenase-1 expression through p38 MAPK/Nrf2 pathway in human dopaminergic cells. Neurochemistry international 2014, 75:79–88.

46. Zhao X, Zou Y, Xu H, Fan L, Guo H, Li X, Li G, Zhang X, Dong M: Gastrodin protect primary cultured rat hippocampal neurons against amyloid-beta peptide-induced neurotoxicity via ERK1/2-Nrf2 pathway. Brain Res 2012, 1482:13–21.

47. Li M, Qian S: Gastrodin Protects Neural Progenitor Cells Against Amyloid β (1-42)-Induced Neurotoxicity and Improves Hippocampal Neurogenesis in Amyloid β (1-42)-Injected Mice. J Mol Neurosci 2016, 60(1):21–32.

48. Yao YY, Bian LG, Yang P, Sui Y, Li R, Chen YL, Sun L, Ai QL, Zhong LM, Lu D: Gastrodin attenuates proliferation and inflammatory responses in activated microglia through Wnt/beta-catenin signaling pathway. Brain research 2019, 1717:190–203.

49. Yang P, Han Y, Gui L, Sun J, Chen YL, Song R, Guo JZ, Xie YN, Lu D, Sun L: Gastrodin attenuation of the inflammatory response in H9c2 cardiomyocytes involves inhibition of NF-κB and MAPKs activation via the phosphatidylinositol 3-kinase signaling. Biochem Pharmacol 2013, 85(8):1124–1133.

50. Liu CM, Tian ZK, Zhang YJ, Ming QL, Ma JQ, Ji LP: Effects of Gastrodin against Lead-Induced Brain Injury in Mice Associated with the Wnt/Nrf2 Pathway. Nutrients 2020, 12(6).

51. Rai SN, Dilnashin H, Birla H, Singh SS, Zahra W, Rathore AS, Singh BK, Singh SP: The Role of PI3K/Akt and ERK in Neurodegenerative Disorders. Neurotoxicity research 2019, 35(3):775–795.

52. Ka M, Condorelli G, Woodgett JR, Kim WY: mTOR regulates brain morphogenesis by mediating GSK3 signaling. Development 2014, 141(21):4076–4086.

53. Pei JJ, Braak E, Braak H, Grundke-Iqbal I, Iqbal K, Winblad B, Cowburn RF: Distribution of active glycogen synthase kinase 3beta (GSK-3beta) in brains staged for Alzheimer disease neurofibrillary changes. Journal of neuropathology and experimental neurology 1999, 58(9):1010–1019.

54. Wang JZ, Wu Q, Smith A, Grundke-Iqbal I, Iqbal K: Tau is phosphorylated by GSK-3 at several sites found in Alzheimer disease and its biological activity markedly inhibited only after it is prephosphorylated by A-kinase. FEBS letters 1998, 436(1):28–34.

55. Tang Z, Guo M, Peng Y, Zhang T, Xiao Y, Ni R, Qi X: Quercetin reduces APP expression, oxidative stress and mitochondrial dysfunction in the N2a/APPswe cells via ERK1/2 and AKT pathways. bioRxiv 2022:2022.2009.2018.508406.

56. Chen Q, Lai C, Chen F, Ding Y, Zhou Y, Su S, Ni R, Tang Z: Emodin Protects SH-SY5Y Cells Against Zinc-Induced Synaptic Impairment and Oxidative Stress Through the ERK1/2 Pathway. Front Pharmacol 2022, 13:821521.

57. Wang L, Tang Z, Deng Y, Peng Y, Xiao Y, Xu J, Ni R, Qi X: Myricetin protected against Aβ oligomer-induced synaptic impairment, mitochondrial function and oxidative stress in SH-SY5Y cells via ERK1/2/GSK-3β pathways. bioRxiv 2023:2023.2001.2012.523781–522023.523701.523712.523781.

58. Lee SE, Kwon D, Shin N, Kong D, Kim NG, Kim HY, Kim MJ, Choi SW, Kang KS: Accumulation of APP-CTF induces mitophagy dysfunction in the iNSCs model of Alzheimer’s disease. Cell Death Discov 2022, 8(1):1.

59. Shi R, Zheng CB, Wang H, Rao Q, Du T, Bai C, Xiao C, Dai Z, Zhang C, Chen C et al: Gastrodin Alleviates Vascular Dementia in a 2-VO-Vascular Dementia Rat Model by Altering Amyloid and Tau Levels. Pharmacology 2020, 105(7-8):386–396.

60. Mishra M, Huang J, Lee YY, Chua DS, Lin X, Hu JM, Heese K: Gastrodia elata modulates amyloid precursor protein cleavage and cognitive functions in mice. Biosci Trends 2011, 5(3):129–138.

61. Jiang Y, Sato Y, Im E, Berg M, Bordi M, Darji S, Kumar A, Mohan PS, Bandyopadhyay U, Diaz A et al: Lysosomal Dysfunction in Down Syndrome Is APP-Dependent and Mediated by APP-βCTF (C99). J Neurosci 2019, 39(27):5255–5268.

62. Xu W, Weissmiller AM, White JA, 2nd, Fang F, Wang X, Wu Y, Pearn ML, Zhao X, Sawa M, Chen S et al: Amyloid precursor protein-mediated endocytic pathway disruption induces axonal dysfunction and neurodegeneration. J Clin Invest 2016, 126(5):1815–1833.

63. Vaillant-Beuchot L, Mary A, Pardossi-Piquard R, Bourgeois A, Lauritzen I, Eysert F, Kinoshita PF, Cazareth J, Badot C, Fragaki K et al: Accumulation of amyloid precursor protein C-terminal fragments triggers mitochondrial structure, function, and mitophagy defects in Alzheimer’s disease models and human brains. Acta Neuropathol 2021, 141(1):39–65.

64. García-Ayllón M-S, Lopez-Font I, Boix CP, Fortea J, Sánchez-Valle R, Lleó A, Molinuevo J-L, Zetterberg H, Blennow K, Sáez-Valero J: C-terminal fragments of the amyloid precursor protein in cerebrospinal fluid as potential biomarkers for Alzheimer disease. Scientific Reports 2017, 7(1):2477.

65. Luo K, Wang Y, Chen W-S, Feng X, Liao Y, Chen S, Liu Y, Liao C, Chen M, Ao L: Treatment Combining Focused Ultrasound with Gastrodin Alleviates Memory Deficit and Neuropathology in an Alzheimer’s Disease-Like Experimental Mouse Model. Neural Plasticity 2022, 2022:5241449.

66. Wang Y, Luo K, Li J, Liao Y, Liao C, Chen WS, Chen M, Ao L: Focused Ultrasound Promotes the Delivery of Gastrodin and Enhances the Protective Effect on Dopaminergic Neurons in a Mouse Model of Parkinson’s Disease. Front Cell Neurosci 2022, 16:884788.

67. Wegmann S, Biernat J, Mandelkow E: A current view on Tau protein phosphorylation in Alzheimer’s disease. Curr Opin Neurobiol 2021, 69:131–138.

68. He J, Li X, Yang S, Li Y, Lin X, Xiu M, Liu Y: Gastrodin extends the lifespan and protects against neurodegeneration in the Drosophila PINK1 model of Parkinson’s disease. Food Funct 2021, 12(17):7816–7824.

69. Mecocci P, Boccardi V, Cecchetti R, Bastiani P, Scamosci M, Ruggiero C, Baroni M: A Long Journey into Aging, Brain Aging, and Alzheimer’s Disease Following the Oxidative Stress Tracks. Journal of Alzheimer’s disease: JAD 2018, 62(3):1319–1335.

70. Muche A, Arendt T, Schliebs R: Oxidative stress affects processing of amyloid precursor protein in vascular endothelial cells. PloS one 2017, 12(6):e0178127.

71. Yun HM, Jin P, Park KR, Hwang J, Jeong HS, Kim EC, Jung JK, Oh KW, Hwang BY, Han SB et al: Thiacremonone Potentiates Anti-Oxidant Effects to Improve Memory Dysfunction in an APP/PS1 Transgenic Mice Model. Molecular neurobiology 2016, 53(4):2409–2420.

72. Niedzielska E, Smaga I, Gawlik M, Moniczewski A, Stankowicz P, Pera J, Filip M: Oxidative Stress in Neurodegenerative Diseases. Molecular neurobiology 2016, 53(6):4094–4125.

73. Petrozziello T, Bordt EA, Mills AN, Kim SE, Sapp E, Devlin BA, Obeng-Marnu AA, Farhan SMK, Amaral AC, Dujardin S et al: Targeting Tau Mitigates Mitochondrial Fragmentation and Oxidative Stress in Amyotrophic Lateral Sclerosis. Molecular neurobiology 2022, 59(1):683–702.

74. Melov S, Adlard PA, Morten K, Johnson F, Golden TR, Hinerfeld D, Schilling B, Mavros C, Masters CL, Volitakis I et al: Mitochondrial oxidative stress causes hyperphosphorylation of tau. PloS one 2007, 2(6):e536.

75. Moreira PI: Alzheimer’s disease and diabetes: an integrative view of the role of mitochondria, oxidative stress, and insulin. Journal of Alzheimer’s disease: JAD 2012, 30 Suppl 2:S199–215.

76. de Oliveira MR, Brasil FB, Fürstenau CR: Evaluation of the Mitochondria-Related Redox and Bioenergetics Effects of Gastrodin in SH-SY5Y Cells Exposed to Hydrogen Peroxide. J Mol Neurosci 2018, 64(2):242–251.

77. de Oliveira MR, de Bittencourt Brasil F, Fürstenau CR: Inhibition of the Nrf2/HO-1 Axis Suppresses the Mitochondria-Related Protection Promoted by Gastrodin in Human Neuroblastoma Cells Exposed to Paraquat. Mol Neurobiol 2019, 56(3):2174–2184.

78. Li Q, Niu C, Zhang X, Dong M: Gastrodin and Isorhynchophylline Synergistically Inhibit MPP(+)-Induced Oxidative Stress in SH-SY5Y Cells by Targeting ERK1/2 and GSK-3beta Pathways: Involvement of Nrf2 Nuclear Translocation. ACS chemical neuroscience 2018, 9(3):482–493.

79. Ting HC, Yang HI, Harn HJ, Chiu IM, Su HL, Li X, Chen MF, Ho TJ, Liu CA, Tsai YJ et al: Coactivation of GSK3β and IGF-1 Attenuates Amyotrophic Lateral Sclerosis Nerve Fiber Cytopathies in SOD1 Mutant Patient-Derived Motor Neurons. Cells 2021, 10(10).

80. Dai JN, Zong Y, Zhong LM, Li YM, Zhang W, Bian LG, Ai QL, Liu YD, Sun J, Lu D: Gastrodin inhibits expression of inducible NO synthase, cyclooxygenase-2 and proinflammatory cytokines in cultured LPS-stimulated microglia via MAPK pathways. PLoS One 2011, 6(7):e21891.

81. Zuo W, Xu F, Zhang K, Zheng L, Zhao J: Proliferation-enhancing effects of gastrodin on RSC96 Schwann cells by regulating ERK1/2 and PI3K signaling pathways. Biomed Pharmacother 2016, 84:747–753.

82. Cruz E, Kumar S, Yuan L, Arikkath J, Batra SK: Intracellular amyloid beta expression leads to dysregulation of the mitogen-activated protein kinase and bone morphogenetic protein-2 signaling axis. PloS one 2018, 13(2):e0191696.

83. Qu LL, Yu B, Li Z, Jiang WX, Jiang JD, Kong WJ: Gastrodin Ameliorates Oxidative Stress and Proinflammatory Response in Nonalcoholic Fatty Liver Disease through the AMPK/Nrf2 Pathway. Phytotherapy research: PTR 2016, 30(3):402–411.

84. Llorens-Martín M, Fuster-Matanzo A, Teixeira CM, Jurado-Arjona J, Ulloa F, Defelipe J, Rábano A, Hernández F, Soriano E, Avila J: GSK-3β overexpression causes reversible alterations on postsynaptic densities and dendritic morphology of hippocampal granule neurons in vivo. Molecular psychiatry 2013, 18(4):451–460.

85. Bradley CA, Peineau S, Taghibiglou C, Nicolas CS, Whitcomb DJ, Bortolotto ZA, Kaang BK, Cho K, Wang YT, Collingridge GL: A pivotal role of GSK-3 in synaptic plasticity. Frontiers in molecular neuroscience 2012, 5:13.

86. Godemann R, Biernat J, Mandelkow E, Mandelkow EM: Phosphorylation of tau protein by recombinant GSK-3beta: pronounced phosphorylation at select Ser/Thr-Pro motifs but no phosphorylation at Ser262 in the repeat domain. FEBS letters 1999, 454(1-2):157–164.

87. Wang Y, Yang R, Gu J, Yin X, Jin N, Xie S, Wang Y, Chang H, Qian W, Shi J et al: Cross talk between PI3K-AKT-GSK-3beta and PP2A pathways determines tau hyperphosphorylation. Neurobiology of aging 2015, 36(1):188–200.

88. Llorens-Martin M, Jurado J, Hernandez F, Avila J: GSK-3beta, a pivotal kinase in Alzheimer disease. Frontiers in molecular neuroscience 2014, 7:46.

89. Sereno L, Coma M, Rodriguez M, Sanchez-Ferrer P, Sanchez MB, Gich I, Agullo JM, Perez M, Avila J, Guardia-Laguarta C et al: A novel GSK-3beta inhibitor reduces Alzheimer’s pathology and rescues neuronal loss in vivo. Neurobiology of disease 2009, 35(3):359–367.

90. Engel T, Hernández F, Avila J, Lucas JJ: Full reversal of Alzheimer’s disease-like phenotype in a mouse model with conditional overexpression of glycogen synthase kinase-3. The Journal of neuroscience: the official journal of the Society for Neuroscience 2006, 26(19):5083–5090.

91. Gourmaud S, Paquet C, Dumurgier J, Pace C, Bouras C, Gray F, Laplanche JL, Meurs EF, Mouton-Liger F, Hugon J: Increased levels of cerebrospinal fluid JNK3 associated with amyloid pathology: links to cognitive decline. J Psychiatry Neurosci 2015, 40(3):151–161.

92. Sclip A, Tozzi A, Abaza A, Cardinetti D, Colombo I, Calabresi P, Salmona M, Welker E, Borsello T: c-Jun N-terminal kinase has a key role in Alzheimer disease synaptic dysfunction in vivo. Cell Death & Disease 2014, 5(1):e1019–e1019.

93. Hoi CP, Ho YP, Baum L, Chow AH: Neuroprotective effect of honokiol and magnolol, compounds from Magnolia officinalis, on beta-amyloid-induced toxicity in PC12 cells. Phytother Res 2010, 24(10):1538–1542.

94. Xiao H, Jiang Q, Qiu H, Wu K, Ma X, Yang J, Cheng O: Gastrodin promotes hippocampal neurogenesis via PDE9-cGMP-PKG pathway in mice following cerebral ischemia. Neurochem Int 2021, 150:105171.

95. Ye T, Meng X, Zhai Y, Xie W, Wang R, Sun G, Sun X: Gastrodin Ameliorates Cognitive Dysfunction in Diabetes Rat Model via the Suppression of Endoplasmic Reticulum Stress and NLRP3 Inflammasome Activation. Front Pharmacol 2018, 9:1346.

