## Supplemental Table 1 for "Gastrodin ameliorates synaptic impairment, reestablishes mitochondrial membrane potential and reduces oxidative stress in N2a/APP cells through ERK1/2 and GSK-3β pathways"

**Supplemental Table 1. Antibodies used in this study.**

| **Antibody** | **Host** | **Dilution** | **Sources** | **Catalog No** |
| --- | --- | --- | --- | --- |
| Anti-Tau S396 | r | 1:1000 | Abcam | #ab32057 |
| Anti-Tau5 | m | 1:1000 | Abcam | #ab80579 |
| SNAP25 | r | 1:5000 | Abcam | #ab5666 |
| Synaptophysin | r | 1:10000 | Abcam | #ab32127 |
| PSD95 | r | 1:2000 | Abcam | #ab18258 |
| Anti-p44/42MAPK(Erk1/2)(137F5) | r | 1:1000 | Cell Signaling | #4695T |
| Anti-P-p44/42MAPK(Erk1/2) (Thr202/Tyr204)(D13.14.4E) | r | 1:2000 | Cell Signaling | #4370S |
| Anti-SAPK/JNK | r | 1:1000 | Cell Signaling | #9252T |
| Anti-p-SAPK/JNK (Thr183/Tyr185) (81E11) | r | 1:1000 | Cell Signaling | #4668S |
| Anti-GSK-3β (D5C5Z) | r | 1:1000 | Cell Signaling | #12456T |
| Anti-p-GSK-3β (Ser9) (5B3) | r | 1:1000 | Cell Signaling | #9323S |
| Anti-APP-CTFs | r | 1:500 | Sigma-Aldrich | #A8717 |
| Anti-β-tubulin | m | 1:5000 | Beyotime | #21463 |
| Anti-GAPDH | r | 1:6000 | Genetex | #GTX100118 |
| Anti-4-Hydroxynonenal | r | 1:200 | Abcam | #ab46545 |
| Anti-8-Hydroxy-2'-deoxyguanosine | m | 1:200 | Abcam | #ab62623 |
| Anti-rabbit IgG (H+L) (HRP) | g | 1:5000 | Thermo Fisher | #31460 |
| Anti-mouse IgG (H+L) (HRP) | g | 1:5000 | Thermo Fisher | #31430 |
| Anti-Mouse IgG (H+L) Alexa Fluor™ 488 | d | 1:200 | Thermo Fisher | #A21202 |
| Anti-Rabbit IgG (H+L) Alexa Fluor™ 546 | d | 1:200 | Thermo Fisher | #A10040 |

APP-CTFs, amyloid precursor protein-derived C-terminal fragments; ERK, extracellular signal-regulated kinase; GAPDH, glyceraldehyde 3-phosphate dehydrogenase; HRP, horseradish peroxidase; d: donkey; g: goat; P: phosphorylated; r: rabbit; m: mouse; T: total; Dilution was used in western blot or immunofluorescence staining.
