## Supplemental Table 2 for "Gastrodin ameliorates synaptic impairment, reestablishes mitochondrial membrane potential and reduces oxidative stress in N2a/APP cells through ERK1/2 and GSK-3β pathways"

**Supplemental Table 2. Chemicals and kits used in this study.**

| **Chemical/kit** | **Sources** | **Catalog No** |
| --- | --- | --- |
| Gastrodin (purity ≥98.0%) | MedChemExpress | #HY-N0115 |
| N2a/APP cell | Tongji Medical School | - |
| Cell Counting Kit-8 | MedChemExpress | #HY-K0301 |
| DAPI Fluoromount-G | Southern Biotech | #0100-20 |
| Dulbecco’s modified Eagle’s medium | Gibco | #11960044 |
| Fetal Bovine Serum | Gibco | #10099141C |
| 5× FD TM DualColor Protein Loading Buffer |  |  |
| Dimethyl sulfoxide | Sigma-Aldrich | #D2650 |
| PEG 300 | Sigma-Aldrich | #91462 |
| Tween-80 | Sigma-Aldrich | #P1754 |
| Triton™ X-100 | Sigma-Aldrich | #X100 |
| Mitochondrial membrane potential assay kit with JC-1 | Beyotime | #C2006 |
| 2’,7’-dichlorofluorescein diacetate assay kit | Beyotime | #S0033S |
| PVDF membranes | Millipore, USA | #ISEQ00010 |
| Non-fat milk | Coolaber, China | #CN7861 |
| Phosphate-buffered saline | Biological Industries, Israel | #02-020-1A |
| SDS polyacrylamide gels | Absin, China | #abs9382 |

DAPI, 4′,6-diamidino-2-phenylindole; DCFH-DA, 2’,7’-dichlorofluorescein diacetate; SDS, Sodium dodecyl sulfate; JC-1, 5,5,6,6’-tetrachloro-1,1’,3,3’-tetraethylbenzimi-dazoylcarbocyanine iodide;
